# Pathway-extended gene expression signatures integrate novel biomarkers that improve predictions of patient responses to kinase inhibitors

**DOI:** 10.1101/2020.11.13.381798

**Authors:** Ashis J. Bagchee-Clark, Eliseos J. Mucaki, Tyson Whitehead, Peter K. Rogan

## Abstract

Cancer chemotherapy responses have been related to multiple pharmacogenetic biomarkers, often for the same drug. This study utilizes machine learning to derive multi-gene expression signatures that predict individual patient responses to specific tyrosine kinase inhibitors, including erlotinib, gefitinib, sorafenib, sunitinib, lapatinib and imatinib. Support Vector Machine learning was used to train mathematical models that distinguished sensitivity from resistance to these drugs using a novel systems biology-based approach. This began with expression of genes previously implicated in specific drug responses, then expanded to evaluate genes whose products were related through biochemical pathways and interactions. Optimal pathway-extended support vector machines predicted responses in patients at accuracies of 70% (imatinib), 71% (lapatinib), 83% (sunitinib), 83% (erlotinib), 88% (sorafenib) and 91% (gefitinib). These best performing pathway-extended models demonstrated improved balance predicting both sensitive and resistant patient categories, with many of these genes having a known role in cancer etiology. Ensemble machine learning-based averaging of multiple pathway-extended models derived for an individual drug increased accuracy to >70% for erlotinib, gefitinib, lapatinib, and sorafenib. Through incorporation of novel cancer biomarkers, machine learning-based pathway-extended signatures display strong efficacy predicting both sensitive and resistant patient responses to chemotherapy.

## 1. Introduction

Selection of a chemotherapy regimen is largely determined by efficacy of a drug in eligible subjects for a specific type and stage of cancer, and considers duration, location and magnitude of responses.^1^ Individuals progress to second-line chemotherapeutic agents after demonstrating or developing limited efficacy to or after relapse from first-line chemotherapeutics.^2,3^ It is feasible to consider personal differences in genomic responses as a means of differentiating between acceptable chemotherapies with otherwise similar response rates across populations of eligible patients.^4^

Previously, we developed gene signatures that predict patient responses to specific chemotherapies from gene expression (GE) and copy number (CN) levels in a set of distinct breast and/or bladder cancer cell lines,^5^ with each line characterized by the drug concentration that inhibited growth by half (GI_50_).^6,7^ Support vector machine (SVM) and random forest machine learning (ML) models were built for each drug using expression and/or copy number values from ‘curated genes’ with evidence from published cancer literature of a contribution to the function or response to said drug in cell lines or patients. This paper develops signatures for tyrosine kinase inhibitors (TKIs),^8^ for which literature on genes associated with response is somewhat more limited.

We developed a novel technique for generating biochemically inspired gene signature models by expanding the pool of genes for ML to include genes both possessing and lacking literature support. The premise for including novel genes or gene products in these models is that these candidates could be related to genes supported by documented evidence through biochemical pathways or interactions that also contribute to drug response. We then compare conventional ML-based gene signatures to corresponding pathway-extended (PE) versions for these TKIs.

Abnormal expression levels or mutations in tyrosine kinases are often causally related to tumour angiogenesis^9^ and metastasis^10^ in certain cancers.^11,12^ TKIs have emerged as effective anti-cancer therapies, owing to their activity by ATP-competitive inhibition of the catalytic binding site of these kinases.^13^ Despite a conserved mechanism of action, sorafenib, sunitinib, erlotinib, gefitinib, imatinib and lapatinib preferentially inhibit different tyrosine kinase targets and exhibit distinct pharmacokinetic profiles.^13–15^ Sorafenib and sunitinib both inhibit VEGFRs, PDGFRs, FLT3R, RET and c-Kit.^15,16^ However, structural differences produce different binding profiles. For example, in binding VEGFR, sorafenib stabilizes the DFG-out inactive conformation of the enzyme, which allows it to bind within an allosteric pocket,^17^ whereas sunitinib binds in and around the ATP-binding region, imparting lower kinase selectivity and faster off-rates.^18^ Similarly, erlotinib and gefitinib are both preferential inhibitors of EGFR, and share analogous chemical structure;^19,20^ but post-absorption, gefitinib is localized to a greater extent in tumour tissue, while erlotinib preferentially accumulates in plasma.^21^ Imatinib is particularly selective for the ABL kinase^8,22,23^ while lapatinib binds to both EGFR and ERBB2.^24^ The specificities of TKIs for different tyrosine kinase targets and the relative activities of those targets in different tumour types largely determine which of these drugs are recommended to treat individual clinical indications. These include renal cell carcinoma (sunitinib, sorafenib), hepatocellular carcinoma (sorafenib), pancreatic cancer (erlotinib), lung cancer (erlotinib, gefitinib), breast cancer (lapatinib) and chronic myelogenous leukemia (imatinib).

Tumor cells can exhibit intrinsic or acquired resistance to chemotherapy. Intrinsic responses refer to an inherent capability to suppress the effects of treatment or render treatment cytostatic to functional characteristics of these cells. In acquired resistance, the tumor mutates or undergoes epigenetic changes after an initial period of clinical success that renders it impervious to treatment.^25,26^ Cytostasis is often achieved by inhibition of glycolytic activity with signal transduction, with the largest group of drugs targeting tyrosine kinases.^27^ On average, tumors initially responsive to TKI treatments such as erlotinib and gefitinib will progress again within a year of treatment.^28,29^ Intrinsic resistance to these TKI drugs tends to be uncommon in EGFR-positive tumors.^30^

Recent studies have revealed novel pathways of resistance and sensitivity to chemotherapeutic drugs.^31,32^ This study aimed to generate models that comprehensively represent global drug responses by inclusion of novel genes or gene products discoverable through their interactions with gene products known to influence these responses. We modify supervised ML-based models to systematically identify novel biomarkers whose expression is related to GI_50_. Gene expression changes in cancer cell lines that expand conventional gene signatures beyond an initial curated set of genes are utilized, including or replacing the initial set with other genes that interact with them. The resulting signatures aim to improve accuracy of prediction of individual patient responses to chemotherapies targeted towards tyrosine kinases.

## 2. Methods

### 2.1 Data and preprocessing of cell line and cancer patient datasets

Microarray GE, CN, and GI_50_ values of breast cancer cell lines treated with erlotinib, gefitinib, imatinib, lapatinib, sorafenib and sunitinib (obtained from Daemen et al. [2013])^5^) were used to derive ML-based gene signatures that predict drug responses. The median GI_50_ values for these cell lines were applied as the threshold distinguishing sensitivity from resistance during ML. The median and range of GI_50_ values for erlotinib was 4.71 [4.18 - 6.54]; gefitinib was 5.03 [4.48 – 6.45]; imatinib was 4.69 [3.82 – 5.81]; sorafenib was 4.27 [3.0 – 5.83]; and sunitinib was 5.23 [4.70 – 5.98]).^5,6^ For lapatinib, the threshold was set at the GI_50_ value with the maximum difference relative to adjacent cell lines (4.94 [ranges from 4.78 to 6.40]), since the GI_50_ of multiple cell lines were equal to the median value.

Performance of these gene signatures was assessed using published studies of cancer patients treated with these drugs. NCBI Gene Expression Omnibus (GEO; https://www.ncbi.nlm.nih.gov/geo/) sourced datasets contained GE data and linked clinical outcomes of each patient with non-small cell lung carcinoma (NSCLC; GSE61676, N=43)^33^ treated with erlotinib [in combination with bevacizumab], hepatocellular carcinoma (GSE109211, N=67)^34^ treated with sorafenib, breast cancer (GSE66399, N=31)^35^ treated with lapatinib [‘Arm B’ patient set only, which received lapatinib in combination with paclitaxel, fluorouracil, epirubicin and cyclophosphamide], chronic myelogenous leukemia (GSE14671, N=23)^36^ treated with imatinib, breast cancer patients (GSE33658, N=11)^37^ treated with gefitinib [in combination with anastrozole and fulvestrant], and gliomas (GSE51305, N=18)^38^ treated with sunitinib. Each of these studies provided clinical information that included a treatment outcome measure that could then be utilized as a binary outcome measure for comparison with predictions made by various models. These outcome measurements vary from study to study. For patients treated with sorafenib or imatinib, a chemotherapy response biomarker was used to distinguish sensitive from resistant patients. For patients treated with erlotinib or lapatinib, outcome (i.e. survival vs death) was used as a surrogate for response. Cancer cell migration data distinguished patients sensitive vs. resistant to sunitinib (where those with ‘moderate induction’ or ‘moderate inhibition’ were defined as resistant, and those with ‘strong inhibition’ were considered sensitive to the drug). Responses to gefitinib were classified based on Response Evaluation Criteria In Solid Tumors (RECIST) guidelines (where those with progressive disease are considered TKI resistant).^39^

Patient selection criteria differed between studies. In the GSE61676 study (erlotinib), patient data was acquired from the SAKK 19/05 trial, where selection criteria consisted of patients with newly diagnosed or recurrent Stage IIIB or Stage IV NSCLC.^33^ In the sorafenib study (GSE109211), tumour tissue was collected from the STORM trial, which enrolled patients with hepatocellular carcinoma with complete radiological response after surgical resection or local ablation.^34^ The lapatinib study (GSE66399) utilized data from the CHER-LOB study, where female adults with HER2+ breast cancer were selected.^35^ In the GSE33658 patient cohort, CD34+ cells were isolated from peripheral blood collected from newly diagnosed chronic-phase chronic myelogenous leukemia patients treated with imatinib.^36^ In the gefitinib study (GSE33658), biopsies were taken from postmenopausal women with newly diagnosed ER+ breast cancer receiving anastrozole, fulvestrant and gefitinib.^37^ In the sunitinib study (GSE51305), native glioma tissue samples were collected from patients with a diagnosis of high-grade glioma WHO (World Health Organization) grade III or IV who underwent surgical resection.^38^

Different expression microarray platforms were used in these GEO datasets, for example, GSE66399, GSE61676 and GSE51305 each measure GE values with distinct vendor and gene sets. To minimize batch effects and apply the cell-line based signatures to these patient datasets, the data were first normalized on a common scale using quantile normalization, according to our previously published approach.^40^ If multiple microarray probes existed for the same gene, the mean of all probe measurements were determined.

### 2.2 Multiple factor analysis and gene set expansion

Genes associated with therapeutic response or function were curated from previous peer-reviewed publications for each TKI (refer to Additional References). Inclusion criteria were based on evidence of the gene or protein contributing to pharmacokinetic or pharmacodynamic response, or were established biomarkers of sensitivity or resistance. Multiple Factor Analysis (MFA) was performed using cell line expression and GI_50_ (concentration of drug inhibiting 50% growth) data^5^ for each curated gene using the *MFAPreselection* software we have developed (available in a Zenodo archive^41^). The archive describes the algorithm used by *MFAPreselection* to traverse pathway networks, dataflow within the program, and software code. MFA determines the relationship between GI_50_ and GE and/or CN data for all expressed genes as an angle that indicates the degree to which expression or copy number correlates either directly (~ zero degrees) or inversely (~180 degrees) with the GI_50_ of the set of cell lines.^42,43^

Circular plots, generated by *MFAPreselection*, indicate this correlation angle (Figure 1).^43^ *MFAPreselection* searches for known gene pseudonyms and substitutes the correct alias (from www.genecards.org [downloaded July 2016]). In the Daemen et al. dataset^5^ used for training in SVM learning (see below), the microarray platform data was in some instances labeled with conflicting gene names. During pathway extension, associated genes were related to older gene aliases that have been deprecated and reassigned by HUGO (Human Genome Organization) Gene Nomenclature Committee to other unrelated genes. This led to some spurious associations between genes during pathway extension. Examples include: *PPY* which was mismatched due to its former alias ‘PNP’ as well as *DDR2* due to incorrect association with its former alias ‘*TKT*’. For the sorafenib model **PE-Sor**, associations of *GC* to *CNN1* and *CA3* were eliminated due to its original designation as ‘DBP’, however its associations between *HNF1A*, *CYP11B1*, *CYP27B1*, and *PIK3R3* remained valid (Figure S1). This issue was addressed using a program script that removed these unsupported associations from the output of *MFAPreselection*.^41^

**Figure 1.**
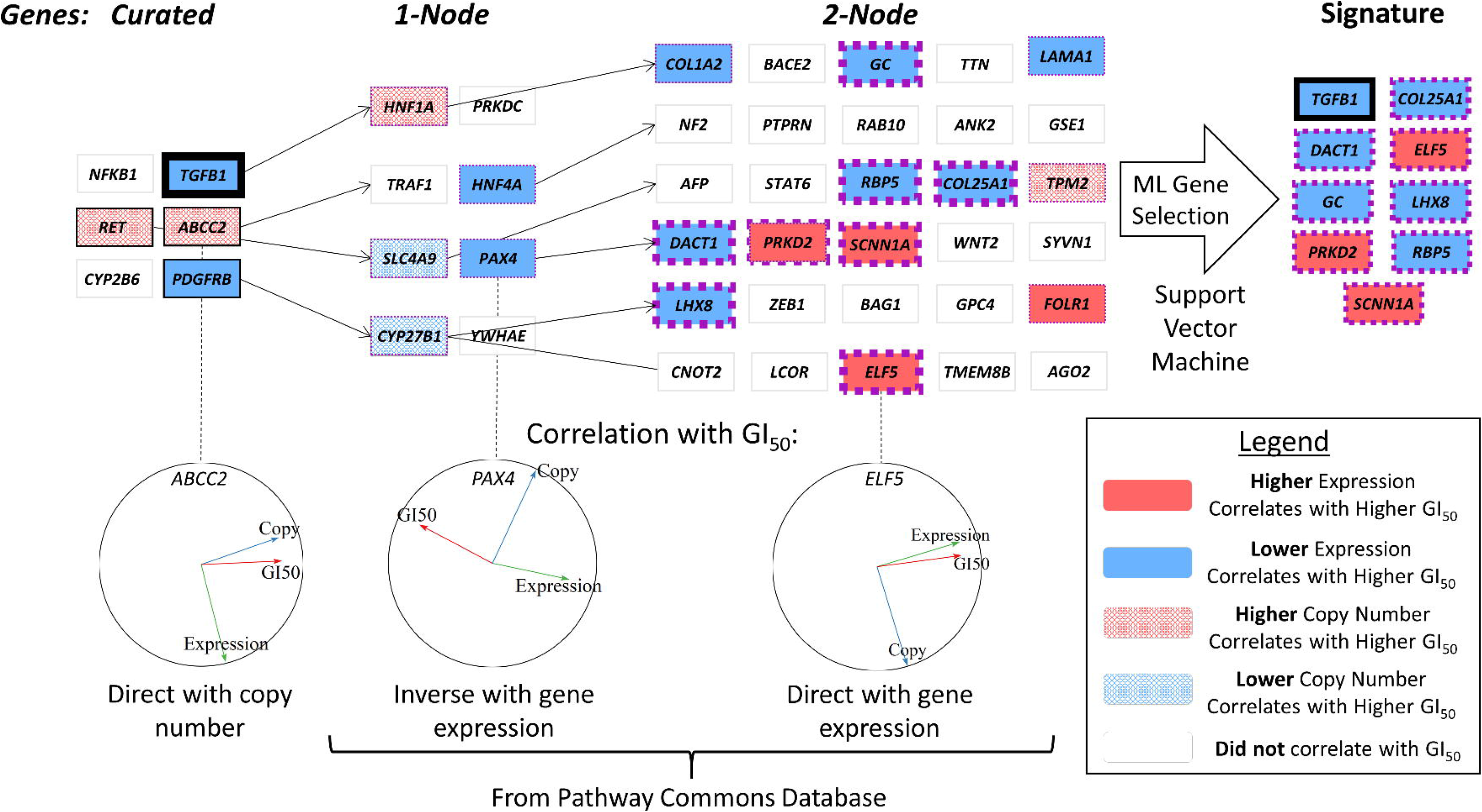
Procedure for Pathway Gene Selection. An initial set of genes with known associations to a particular TKI (here we show a subset of sorafenib-related genes) are selected and then evaluated by MFA, which was used to find a correlation between cell line drug sensitivity (GI_50_) and the GE or CN of these genes in those cell lines (*left*). MFA correlation circles visualize these relationships (*bottom*). The gene list is extended, using pathway and interaction databases (i.e. PathwayCommons) to find genes related to curated genes which showed MFA correlation to GI_50_ (one-node distant genes; *middle-left*). The list is extended again from the MFA-correlating one-node distant genes (two-node distant genes; *middle-right*). All curated and extended genes which showed an MFA correlation were then used as features to generate a final predictive SVM gene signature for the evaluated TKI (*right*). Genes within the best performing sorafenib signature are indicated in thick borders (black for curated genes, purple for pathway-extended genes).

A Perl script was written to eliminate these spurious matches by confirming relations reported by *MFAPreselection* with the PathwayCommons Interaction SIF (Simple Interaction Format) file (“Parentage-MFA-Path-Source-Program.Simple-Output-Version.pl”; provided in a Zenodo archive^41^). If corrected labels were not found or a gene was absent from a microarray platform, then this cell line or gene is not included in the analysis.

ML signatures were expanded by *MFAPreselection* to include genes associated with curated genes by extension using components of adjacent biochemical pathways (pathway-extension, or PE; Figure S2). To identify these relationships, *MFAPreselection* relied on the PathwayCommons database (version 8 [downloaded April 2016]) to assess expanded gene lists by inclusion of genes addressable from the curated set (one node distant from a curated gene), followed by a second iteration (two nodes distant from a curated gene; illustrated in Figure 1). During this process, genes that did not meet minimally-defined levels of MFA correlation to drug GI_50_ (either positive or negative) were discarded and additional gene expansion steps also ignored these genes. These levels were determined using six different conditions set for the *MFAPreselection* software: maximum thresholds up to 10° and 20° from either full direct or inverse correlation for curated genes only (conditions #1 and 2, respectively); up to a 10° and 20° threshold for both curated genes and directly related genes (one-node distant; conditions #3 and 4, respectively); and up to a 10° and 20° threshold for curated genes and genes up to two nodes distant from the curated gene set (conditions #5 and 6, respectively). Genes in which GI_50_ was correlated with CN (Tables S1 [A-F]) were not considered for SVM analyses due to unavailability of CN data in patient datasets.

### 2.3 SVM learning

Genes with expression levels correlated with GI_50_ were qualified for SVM analysis. SVMs were used to train GE datasets against GI_50_ data using the MATLAB statistics toolbox (similar to the procedure described in Mucaki et al. [2016]^44^ using SVM software developed in Zhao et al. [2018]^40^; software available at: doi:10.5281/zenodo.1170572). Instead of using the “*fitcsvm*” function (as in Mucaki et al. [2016]^44^), a multiclass-compatible “*fitcecoc*” function was used to generate SVM signatures, with both misclassification rate^44^ and log loss^40^ value used as performance metrics to derive optimal signature models. A forward feature selection (FFS) algorithm was used to generate these gene signatures (program from Zhao et al. [2018]^40^: “*FFS_strat_kfold_gridsearch.m”*). FFS tests each gene at random from the qualified gene set by training a cross-validated Gaussian kernel SVM on the training data to determine the individual gene that produces the lowest misclassification rate or log loss value. Subsequent genes are then added to determine whether model performance is improved, until the performance criterion converges to a minimum value. Models were built using a range of C and sigma values (from 1 to 100,000, in multiples of 10 for each variable [where C ≥ sigma]; 21 total combinations). Since the goal of pathway extension was to expand and improve these models beyond curated signatures with ≥ 2 genes, PE-derived gene signatures with fewer than two genes were excluded from proceeding to the validation step.

### 2.4 Validation of Cell-Line Derived Gene Signatures using Patient Data

All derived multi-gene SVMs were validated against clinical patient data using traditional validation (MatLab program “*regularValidation_multiclassSVM.m*’ from Zhao et al. [2018]^40^). Performance was indicated by both overall predictive accuracy and by Matthews Correlation Coefficient (MCC, which assesses overall quality of a binary classifier by considering the balance of true and false positives and negatives). Overall, the best-performing gene signature for each drug was selected by MCC, as it is a metric not skewed by imbalanced data. Once the best performing SVM for each drug was established, leave-one-out cross-validation^7^ was used to determine the overall impact of each individual gene to the model itself (change in misclassification or log loss), as well as its impact on the accuracy of the model to predict chemotherapy response. Top-performing PE TKI models can be accessed to predict responses based on expression in individual patients with our web-based SVM calculator (http://chemotherapy.cytognomix.com).^6^

Ensemble averaging of multiple SVM models involved weighting patient predictions from highest performing models derived for a particular TKI by the area under the curve (AUC) of each corresponding model (computed using the MATLAB function ‘*perfcurve*’). MCC itself was also evaluated as a potential source of weights for ensemble averaging; however, AUC-weighted predictions were superior in overall performance. The number of models included in the ensemble varied, as the number of highest performance models for each TKI differed (4 for sorafenib; 2 for erlotinib, sorafenib, imatinib, sunitinib and gefitinib). A patient was considered resistant to a drug if the sum of all AUC-weighted predictions were > 0 and sensitive if this sum was < 0.

## 3. Results

### 3.1 Generating SVM signatures using breast cancer cell line-training data

Genes associated with drug response or function were curated for gefitinib (N=113), sunitinib (N=90), erlotinib (N=71), imatinib (N=157), sorafenib (N=73), and lapatinib (N=91) (curated genes are provided in Table S1 and labeled as ‘0’ node distant genes). In general, MFA was performed using 48 breast cancer cell lines using GE, CN and GI_50_ values for each gene.^5^ Biochemically inspired ML-based signatures for each TKI, derived from curated genes, were obtained according to our previously described approach.^6^ MFA analysis was also performed on genes encoding proteins related to these curated genes (through interaction or as neighbours in the same biochemical pathway) to identify those that also correlated, either directly or inversely, with GI_50_ (all GI_50_-correlated PE genes are provided in Table S1 [labeled as 1-node and 2-node distant genes]). This expanded set of GI_50_-correlating genes were then used to derive SVMs containing combinations of curated and PE genes. The derived signatures for each TKI minimized either misclassification or log loss to generate the best performing models. The best performing curated and PE SVM signatures for erlotinib [**C-Erl**, **PE-Erl**], sorafenib [**C-Sor, PE-Sor**], gefitinib [**C-Gef, PE-Gef**], lapatinib [**C-Lap**, **PE-Lap**], imatinib [**C-Ima, PE-Ima**], and sunitinib [**C-Sun**, **PE-Sun**] are summarized in Table 1, whereas the performance of all models is indicated in Table S2.

**Table 1.** Performance of Curated SVMs and PE Models on Training and Patient Testing Data. Curated and PE SVMs were derived for each TKI based on ability sorting cancer cell lines. The C (box-constraint), σ (kernel-scale) and features comprising the best-performing model are indicated. Models listed are those which exhibited optimal performance, defined as the model with the highest MCC against the patient data set. ‘Validation’ indicates the predicted drug response of patients made by each curated and PE model as compared to the observed response provided by these studies. ‘Training’ indicates either percent misclassification or overall log-loss of the cell line-based model by cross-validation, depending on which minimization metric was used in said model derivation.

| TKI<br>(Patient Tumor Type;<br>GEO Dataset) | Model | Gene Signature (SVM: C; $\sigma$) | Validation | | | | Training | |
| --- | --- | --- | --- | --- | --- | --- | --- | --- |
|  |  |  | MCC | Sensitive | Resistant | Overall | Log Loss | Misclassification |
| **Erlotinib**<br>(NSC Lung<br>Carcinoma;<br>GSE61676) | Cur | *BAX, FOXO1* (100; 1) | 0.08 | 100% | 3% | 23% | - | 0.38 |
|  | PE | *NEK7, SLCO3A1, RELB, FRMD4A, HSD17B2, CDK6, PALM, IL1RN, SMYD1, BAG2, GNG3, SULT1E1* (1000; 100) | 0.41 | 42% | 93% | 83% | 0.01 | - |
| **Sorafenib**<br>(Hepatocellular<br>Carcinoma;<br>GSE109211) | Cur | *PDGFRB, TGFB1, SLCO1B1* (10000; 1) | 0.28 | 96% | 29% | 50% | 0.72 | - |
|  | PE | *ELF5, RBP5, GC, PRKD2, SCNN1A, COL25A1, TGFB1, DACT1, LHX8* (100000; 1000) | 0.72 | 72% | 95% | 88% | 0.05 | - |
| **Gefitinib**<br>(Breast Cancer;<br>GSE33658) | Cur | *GRP* (10000; 10) | 0.16 | 13% | 100% | 29% | 0.53 | - |
|  | PE | *CNTN1, CXCL2, NTRK3, GCG* (10000; 10) | 0.67 | 100% | 50% | 91% | 0.58 | - |
| **Lapatinib**<br>(Breast Cancer;<br>GSE66399) | Cur | *ERBB2* (10000; 1) | 0.31 | 13% | 100% | 77% | 0.72 | - |
|  | PE | *FBP1, ITGA11, TRIM68, BCAT1, ZNF780A, UTP20, GRB7* (10; 10) | 0.33 | 63% | 74% | 71% | 0.01 | - |
| **Imatinib**<br>(Leukemia;<br>GSE14671) | Cur | *IL3, ABL2, CDKN1A* (10000; 10) | 0.23 | 41% | 83% | 52% | - | 0.42 |
|  | PE | *LIF, MRGPRF, GRM3, TNNI1, CACNA2D1* (100000; 100) | 0.18 | 84% | 33% | 70% | - | 0.06 |
| **Sunitinib**<br>(Glioma;<br>GSE51305) | Cur | *HGF, VEGFC, TSC1, AXL, ENPP2, NFKB1* (100000; 10000) | 0.31 | 87% | 45% | 59% | - | 0.14 |
|  | PE | *EPHA2, NR4A1, SIAE* (100000; 100000) | 0.61 | 67% | 92% | 83% | - | 0.20 |

Cur - Curated models derived from genes associated with drug in literature; PE - Pathway-extended models; MCC - Matthews Correlation Coefficient; Sensitive - Accuracy to Drug Responsive Patients; Resistant - Accuracy to Non-Responsive Patients; Overall - Combined accuracy

### 3.2 Validation of cell line-based SVM signatures using cancer patient data

Cell line-derived SVMs for TKIs were initially evaluated on patient data sets where patients were treated with the same agent.^40^ Erlotinib signatures were validated using patients with NSCLC (GSE61676; N=9 survived, 34 died), sorafenib signatures were validated using patients with hepatocellular carcinoma (GSE109211; N=21 sensitive, 46 resistant), sunitinib signatures were validated using outcomes of patients with high-grade gliomas (GSE51305; N=6 sensitive, 12 resistant), imatinib signatures were validated using outcomes of patients with chronic myelogenous leukemia (GSE14671; N=17 sensitive, 6 resistant), and lapatinib and gefitinib signatures were validated based on breast cancer outcomes (GSE66399 [N=8 survived, 23 died] and GSE33658 [N=10 sensitive, 2 with resistant], respectively).

MCC (range −1 to +1) was the primary determinant of model performance, as it measures overall accuracy (OA) while accounting for representation between binary prediction categories.^45^ This was necessary, as patient datasets available exhibited imbalances in the ratios of responsive to non-responsive patients in terms of their respective observed clinical outcomes. Models based on features generated under relaxed constraints (condition #6) generated the best performing SVM on patient data for every TKI, except sorafenib. The best-performing PE model was **PE-Sor**, which accurately predicted patient responses with 0.72 MCC (and 88% OA). The best performing curated model was **Cur-Lap**, with 0.31 MCC (and 77% OA). In comparison to curated SVMs, PE SVMs predicted patient response with 0.26 higher MCC and 33% higher OA (13% increase in accuracy predicting sensitive patients; 13% increase in accurately predicting resistant patients). Except for imatinib, the best-performing PE model outperformed their curated counterpart. This difference in performance is evident in Figure 2, as predictive accuracy for PE models is consistently higher for both resistant and sensitive patient outcomes.

**Figure 2.**
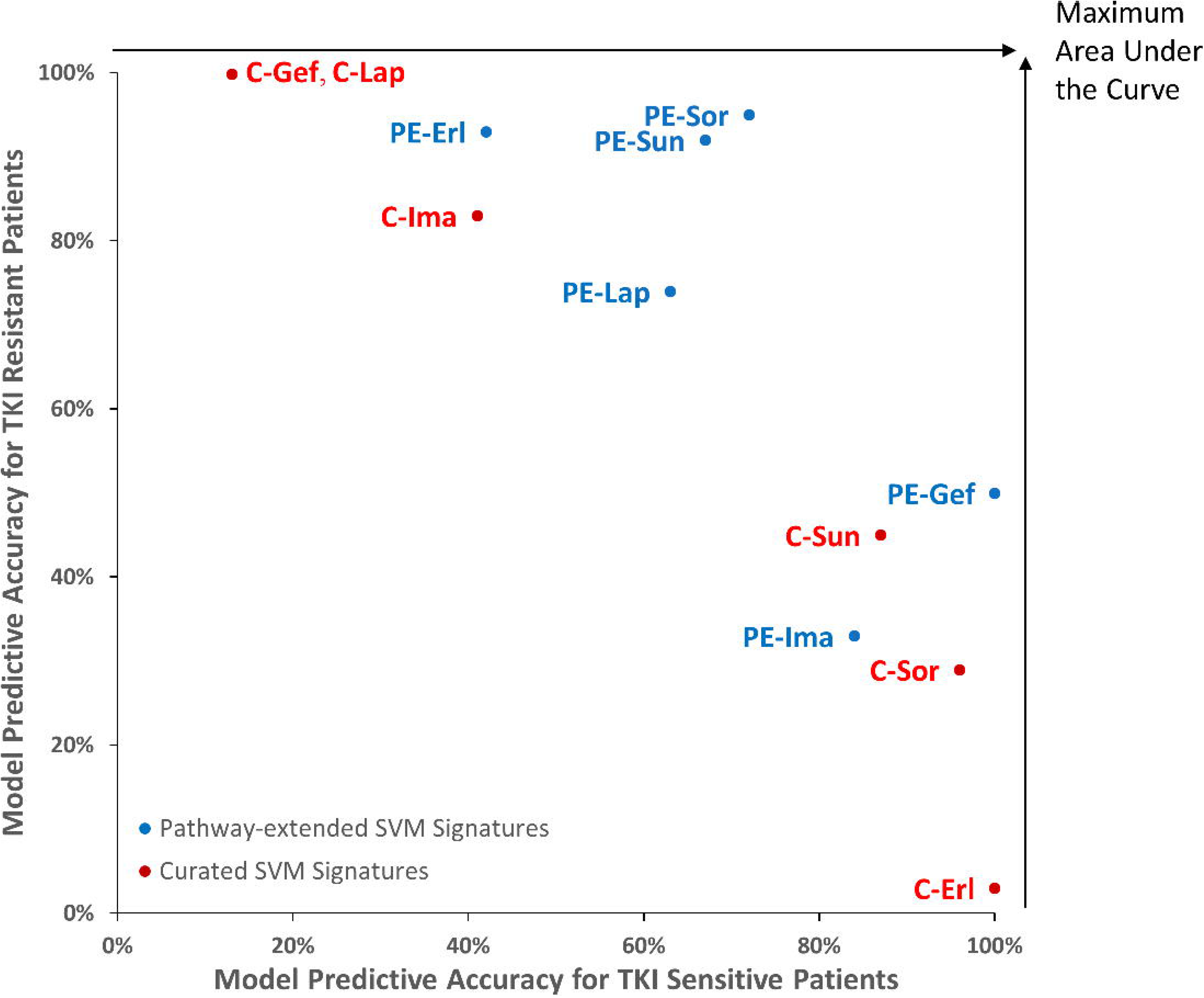
Accuracy of Curated and Pathway-Extended SVMs on TKI Sensitive and Resistant Patients. The predictive accuracy of the best-performing curated (C-) and Pathway-Extended (PE-) models for each TKI were arranged based on their accuracy in classification of drug sensitive and resistant tumour patients. This illustrates how curated models are often only accurate towards one patient class (sensitive or resistant) but not both (red), which is an issue as the patient data was often imbalanced (number of sensitive | resistant patients in each study: lapatinib [‘Lap’; n= 8 | 23], imatinib [‘Ima’; n= 17 | 6], sunitinib [‘Sun’; n= 6 | 12], erlotinib [‘Erl’; n= 9 | 34], gefitinib [‘Gef’; n= 10 | 2], and sorafenib [‘Sor’; n= 21 | 46]). Conversely, predictions by PE SVMs were often more balanced (blue), possessing moderate to high accuracy for both sensitive and resistant patients, and consequently greater accuracy as a whole.

The erlotinib (GSE61676) and gefitinib (GSE33658) studies utilized for model testing provide patient GE data both pre- and post-treatment. This provided an opportunity to determine whether to determine whether short term drug exposure altered GE and model accuracy. For erlotinib, blood samples were obtained prior to and 24 hours post-treatment. For gefitinib, biopsies were taken prior to and 3 weeks post-treatment. Both **PE-Erl** and **PE-Gef** exhibited slightly lower performance for the pre-treatment samples (Table S3), with 5 additional patients misclassified with **PE-Erl** (73% OA with N=43 total patients) and 2 additional misclassified individuals with **PE-Gef** (73% OA; N=12 patients). MCC for **PE-Gef** is significantly lower (−0.15), since the model misclassifies all untreated individuals as resistant. Treatment with these drugs perturbs predictions, but to a limited extent.

### 3.3 Composition of PE SVM signatures and contributions of individual features

PE SVM signatures contain either genes from peer-reviewed literature about the drug response (“initial” or “curated” genes), those related to these genes through direct interactions or as neighbours within the same pathways (one-node distant genes), or genes associated with these one-node distant genes (two-node distant genes). To better comprehend the composition of and relationships between genes in the best-performing PE SVM signatures, we analyzed the connection networks for each model (see Table S4 for connection network for all other top performing PE models). For example, while **PE-Sor** consists of one curated gene and eight two-node distant genes, there are an additional 6 curated and 10 one-node genes that connect the genes in **PE-Sor** by pathway-extension (Figure 3A shows a two-dimensional visualized connection network for this drug; see Figures 3B-F for lapatinib, gefitinib, sunitinib, imatinib and erlotinib, respectively). Due to the complexity of the relationships between gene products for erlotinib, it was not feasible to create an unequivocal two-dimensional network diagram for this drug response, and is instead presented in tabular form (Figure 3F). Nevertheless, it is apparent from the majority of these network diagrams that genes that were two-nodes distant from the curated gene set were most commonly selected in the best performing PE models. Furthermore, the two-node distant genes selected interacted with multiple curated or one-node distant genes.

**Figure 3.**
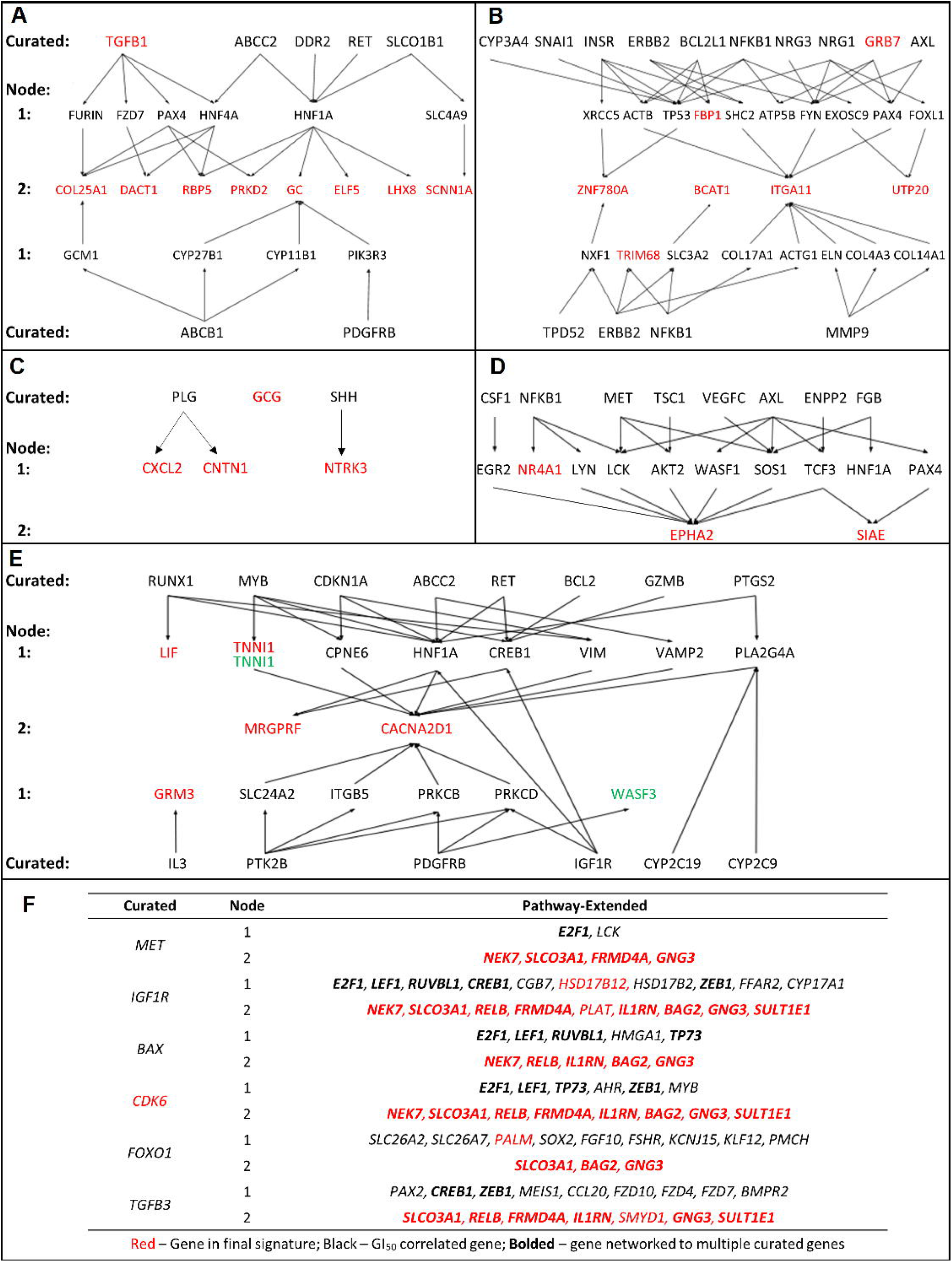
Connection Network for Pathway-Extended TKI SVMs. Schematic relationships outlining the pathway connections for the best-performing PE model for each drug in panels A) sorafenib; B) lapatinib; C) gefitinib; D) sunitinib; E) imatinib and F) erlotinib. All symbols indicated are gene names. The erlotinib model was highly interconnected and is represented as a table. Genes in red are features selected for the final **PE-Sor** gene signature, while genes colored green were chosen in a separate PE gene signature with comparable performance. Genes in black were not part of the final signature themselves but correlated with efficacy to sorafenib by MFA and expanded the gene pool through biochemical connections they possessed to one-node or two-node distant genes.

To determine the degree to which each gene in a signature contributed to the accuracy of the overall model prediction, we performed leave-one-out cross-validation for each gene in the best-performing model for each drug. We then reassessed the predictions of the resultant signature for the observed responses in the cell lines used for model training (Table S5) and for the patient data used for testing (Figure 4). Based on patient data, the gene features eliminated from models that had the highest impact on performance were: *CDK6*, *BAG2*, *SULT1E1*, and *IL1RN* (**PE-Erl**); *CNTN1, GCG* and *NTRK3* (**PE-Gef**); *GRB7* and *BCAT* (**PE-Lap**); *ELF5*, *TGFB1*, *PRKD2*, *RBP5*, and *GC* (**PE-Sor**); *EPHA2* and *SIAE* (**PE-Sun**); and *CACNA2D1* and *GRM3* (**PE-Ima**). Genes removed that improved predictive performance on patient data included *FBP1* (**PE-Lap**), *PLAT* (**PE-Sor**) and *LHX8* (**PE-Sor**).

**Figure 4.**
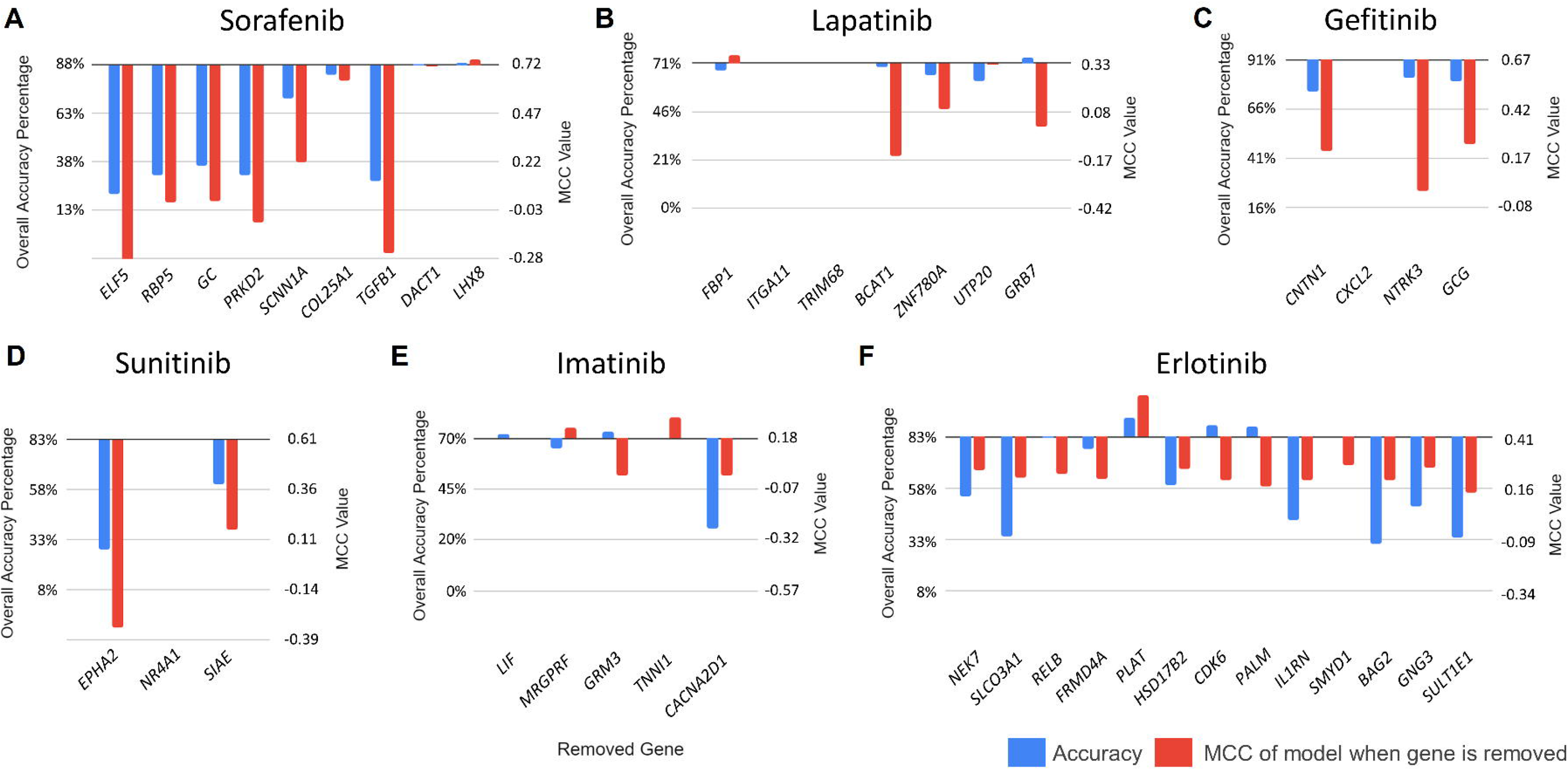
Effect of Removal of Individual Genes from Signature on Overall Accuracy using Patient Tumour Data. The patient classification accuracy and MCC of the strongest performing PE models are altered upon the removal of each component gene listed. These PE TKI gene signatures are: A) sorafenib [**PE-Sor**]; B) lapatinib [**PE-Lap**]; C) gefitinib [**PE-Gef**]; D) sunitinib [**PE-Sun**]; E) imatinib [**PE-Ima**]; and F) erlotinib [**PE-Erl**]. Blue and red bars denote the overall accuracy and MCC of the model after gene removal, respectively.

**PE-Gef** consists of 4 pathway-extended genes (*CNTN1, CXCL2, NTRK3* and *GCG*) and one curated gene, *GCG*. *GCG* encodes a hormone preprotein which is cleaved into four peptides, including glucagon-like peptide 2, which has been found to reduce gefitinib-induced intestinal atrophy in mice.^46^ Removal of *NTRK3* from **PE-Gef** had the largest impact on model performance, reducing MCC to 0. *NTRK3* has a critical role in secretory breast cancer gene, with the *EVT6-NTRK3* fusion oncogene being considered a primary initiating event.^47,48^

**PE-Sun**, which consists of three pathway-extended genes, *SIAE*, *NR4A1, and EPHA2*, was evaluated in gliomas. *NR4A1* is essential for colony formation of glioblastoma cells on soft agar.^49^ Of 14 glioblastoma specimens, 13 possessed elevated *EPHA2* levels.^50^ Removal of *NR4A1* from **PE-Sun** did not alter overall accuracy or MCC of the model, while removal of *EPHA2* decreased overall accuracy by 55% and MCC by 0.94. Regarding *SIAE,* alterations in cell surface sialylation by glucocorticosteroids has been suggested to promote glioma formation.^51^

**PE-Sor** (*COL25A1, TGFB1, DACT1, RBP5, PRKD2, GC, ELF5, LHX8,* and *SCNN1A*) was used to predict sorafenib response in hepatocellular carcinoma (HCC) patients. Removal of *RBP5*, *PRKD2, GC* and *ELF5* significantly reduced overall accuracy (>50%) and MCC (>0.7) (Figure 4A)*. RBP5* is linked to aggressive tumour features in HCC,^52^ *PRKD2* is upregulated in HCC and correlated with metastasis,^53^ and decreased actin-free GC levels have been found to relate with disease severity in HCC.^54,55^ Vitamin D_3_, which is bound by GC, lowers the effective dose of sorafenib required for its cytostatic effect in melanoma and differentiated thyroid carcinoma.^56^ *ELF5* has not been direct connected to HCC, but has been associated with a wide range of cancers.^57,58^ Genes in **PE-Sor** that have not been as strongly linked to cancer (*COL25A1* and *LHX8*) did not change model accuracy to the same extent (<10%) when removed (Figure 4A). Removal of the curated gene *TGFB1*, which enhances the apoptotic activity and sensitizes cells to sorafenib^59^ decreased overall accuracy by 60% in HCC patients. The respective contexts of the curated Sorafenib-related genes juxtaposed with the PE genes in **PE-Sor** are indicated in a cellular schematic of the roles and functions of these genes (Figure 5).

**Figure 5.**
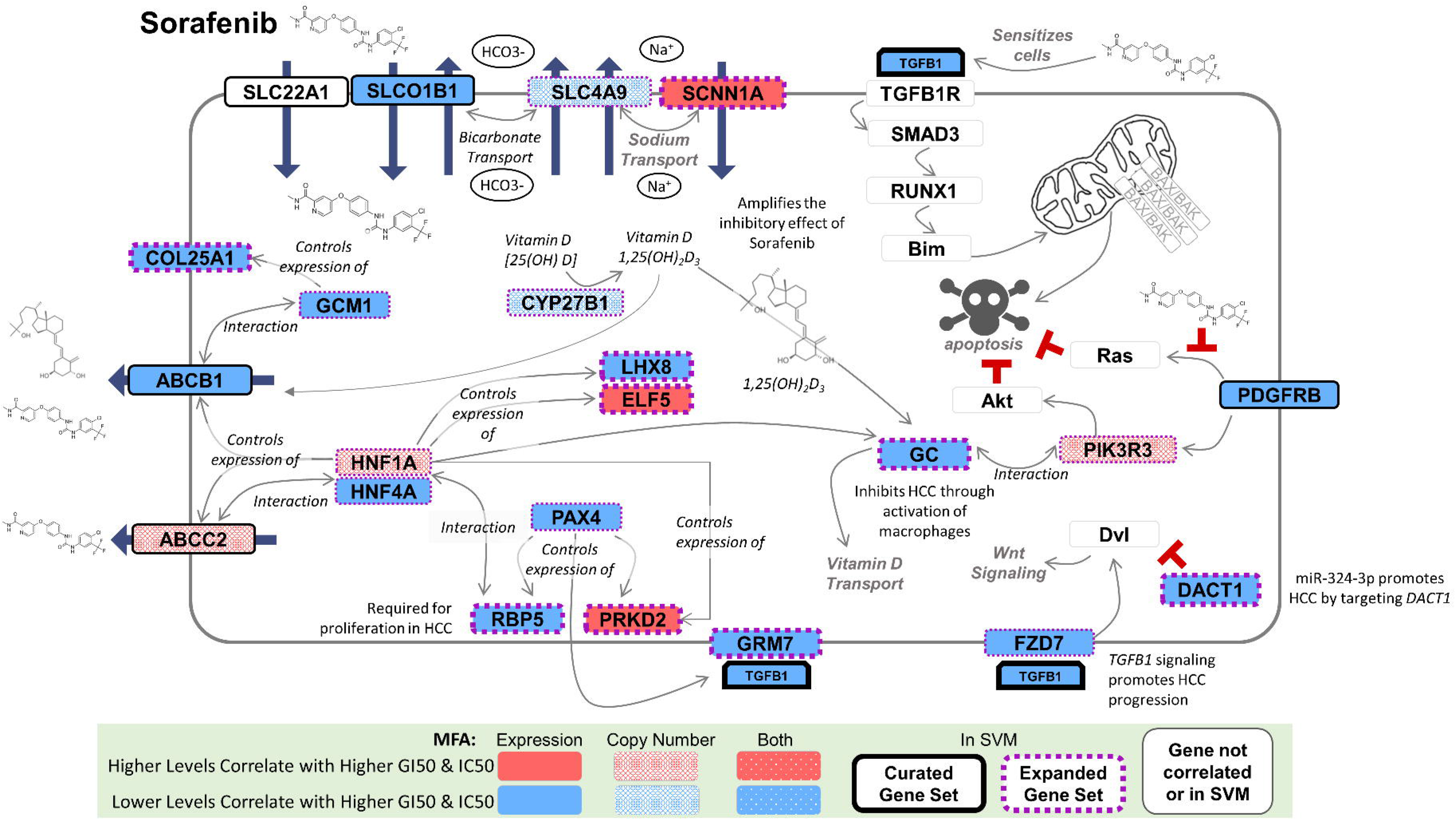
Schematic of the Pathway-Extended Genes in the Sorafenib Model PE-Sor. The best performing sorafenib model **PE-Sor** is a 9-gene model consists of a single curated gene (*TGFB1*) and eight genes selected by pathway-extension (*ELF5*, *RBP5*, *GC*, *PRKD2*, *SCNN1A*, *COL25A1*, *DACT1*, and *LHX8*). This cell schematic provides context of the cellular mechanisms of action and/or known relationships between genes with a documented impact on sorafenib activity (‘curated’ genes; black borders) and those genes selected by pathway extension (purple borders). Genes with grey borders are not curated nor pathway-extended genes and are simply present to give context between genes and their known cellular functions. Thicker borders specify those genes in the **PE-Sor** model, while gene colour-coding indicates how GE and/or copy number correlated to sorafenib GI_50_ by MFA.

**PE-Ima** (*LIF*, *MRGPRF*, *GRM3*, *TNNI1*, and *CACNA2D1*) predicted imatinib response in chronic myelogenous leukemia patients. *LIF* encodes a protein which prevents continued growth of myeloid leukemia cells by inducing terminal differentiation,^60^ although independent removal of *LIF* did not notably affect model performance. Downregulation of *CACNA2D1* from **PE-Ima** is associated with erythroid differentiation of K562 and KCL-22 chronic myeloid leukemia cells.^61^ Removal of *CACNA2D1* decreased both classification accuracy and MCC (−44% and −0.18, respectively; Figure 4E).

A second PE model (indicated in green in Figure 4E) exhibited comparable performance to **PE-Ima**: *TNNI1* and *WASF3* [C=10000, σ=10000], with an OA of 57% (47% accurate with sensitive and 83% with resistant patients; MCC = 0.27). *WASF3* has been implicated in breast cancer metastasis.^62^ *TNNI1*, a gene that is shared by both this model and **PE-Ima**, is one of the three inhibitory subunits of smooth muscle troponin, that are all overexpressed in breast cancer.^63^ Interestingly, the kinase, *TNNI3K*, that phosphorylates this protein is essential for proliferation of mononuclear diploid cardiomyocytes during heart muscle repair due to injury.^64^ Phosphorylation of troponin would appear to have a previously uncharacterized moonlighting function in tumor development.^65^ If imatinib inhibits *TNNI3K* through an off-target effect, this may modulate *TNNI1* activation and possibly, an associated proliferative phenotype.

**PE-Lap** (*FBP1, ITGA11, TRIM68, BCAT1, ZNF780A, UTP20,* and *GRB7*) predicted outcomes of breast cancer patients treated with lapatinib. Independent removal of *BCAT1* reduced accuracy in predicting sensitive patients. Silencing or knockdown of *BCAT1* has been associated with reduced growth of triple negative breast cancer.^66^ Removal of *ITGA11* or *TRIM68* did not alter **PE-Lap** accuracy (Figure 4B).

**PE-Erl** consisted of *NEK7*, *SLCO3A1*, *RELB*, *FRMD4A*, *HSD17B2*, *CDK6*, *PALM*, *IL1RN*, *SMYD1*, *BAG2*, *GNG3,* and *SULT1E1,* and was used to predict chemotherapy response in NSCLC patients. *BAG2* and *SULT1E1* are novel biomarkers of erlotinib efficacy, as removal of either gene led to imbalanced predictions of sensitive patients by this signature. Overexpression of *BAG2* has been associated with poor disease-specific survival in lung cancer,^67^ while the *SULT1E1* polymorphism rs4149525 has been associated with shortened overall survival in NSCLC.^68^ This model originally contained *PLAT*, which when eliminated from the erlotinib dataset of 43 patients significantly increased in overall accuracy (+10%) and MCC (+0.21) of the model predictions (Figure 4F). *PLAT* was therefore considered a false positive result from ML, and therefore eliminated from gene signature. Our post-hoc analysis demonstrated that the majority of genes (75%) in **PE-Erl** were associated with the NSCLC phenotype.

### 3.4 Performance of PE SVM signatures on sex-stratified patients

Previous studies have suggested that females may be more sensitive to TKI treatment than males.^69,70^ We therefore stratified TKI model performance by sex in the GSE61676 data set, which provided patient sex information along with response (19 male [3 sensitive] and 24 female [6 sensitive] patients). Considering all patients, **PE-Erl** predicted patient response with an MCC of 0.41 and 83% overall accuracy (42% and 93% accurate in patients sensitive and resistant to this drug, respectively). In males alone, **PE-Erl**’s overall accuracy was lower (76%), with MCC notably decreased to 0.11, as **PE-Erl** did not predict individuals who were sensitive or resistant to the drug as accurately (27% and 85%, respectively). In females, **PE-Erl** performed better than for the full data set, with 85% OA (MCC = 0.56), of which resistance was predicted with 99% accuracy and sensitivity was predicted with 42% accuracy (Table S6). This indicates that the **PE-Erl** signature more precisely captures factors that contribute to greater sensitivity in females.

The predictive performance of erlotinib PE model **PE-Erl** to the NSCLC dataset GSE61676 was higher in female patients than male patients (0.45 greater MCC; 9% greater OA). This was consistent with the possibility that **PE-Erl** contains gene(s) distinguishing sex-differentiated sensitivity to the drug. Of the 12 genes comprising **PE-Erl**, independent removal of *RELB* and *CDK6* features from the model notably reduced accuracy of the predicted response in female patients that were sensitive to the drug. *RELB* has previously been identified as a sex-discriminatory candidate gene in trichostatin A-treated chronic lymphocytic leukemia cells due to repressed expression in resistant male cells, but upregulation in resistant female cells.^71^ *RELB* also possesses pro-survival functions across multiple cancer types^72–74^ and has been identified as a prognostic biomarker for NSCLC patients.^75^ Overall, *RELB* is a top candidate gene to explain the improved accuracy of **PE-Erl** in female NSCLC patients.

### 3.5 AUC-weighted ensemble model predictions

Ensemble learning consolidates hypotheses of multiple models to potentially improve predictive performance.^76^ For ensemble learning, each model’s AUC was computed and used to weigh predictions made for each model within the ensemble.^77^ There were 4 SVMs for sorafenib possessing strong predictive accuracy with patient derived expression data. Therefore, all were used for ensemble averaging. For the other TKIs, ensemble learning combined the top- and second-best performing SVMs (Table 2). Ensemble averaging improved both OA and MCC for erlotinib (OA: 84% [+1%]; MCC – 0.45 [+0.04]), and sorafenib (OA: 91% [+3%]; MCC: 0.79 [+0.07]). For patients with the same predicted outcome in ≥75% of cases after ensemble learning, overall accuracy exceeded 80% for all TKIs except lapatinib. Discordant consensus predictions between multiple signatures for the same drug (majority outcome occurred <75% for each patient) exhibited lower overall accuracy.

**Table 2.**
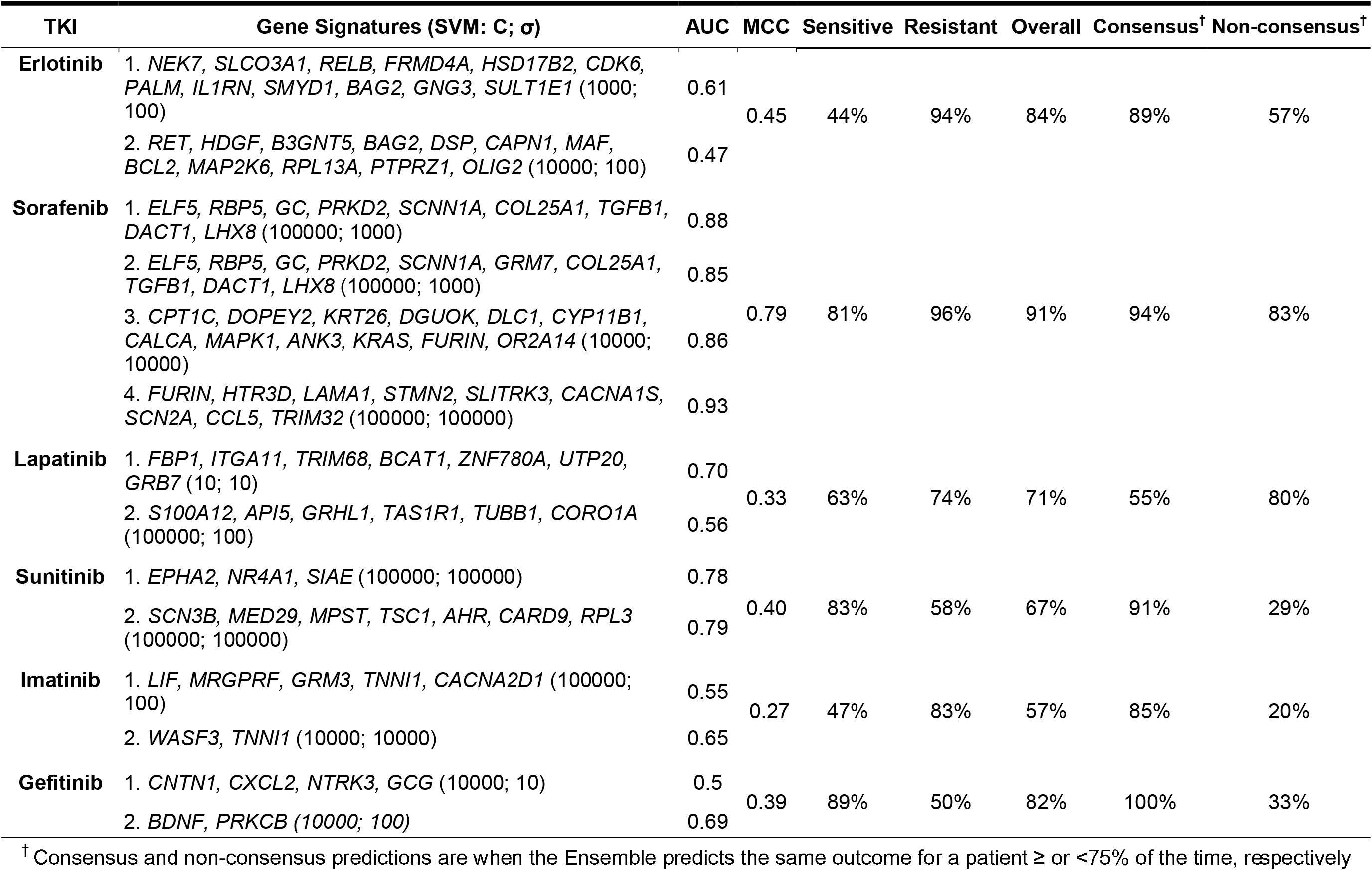
Models used in the Ensemble Averaging analysis of Patient Data. Ensemble averaging amalgamates predictions from numerous SVMs for an individual TKI, weighted by AUC (indicated). Each SVM signature included within ensemble averaging predicted the response of each patient treated with its associated TKI, and the majority prediction was used of that of the ensemble. Overall, the accuracy of the ensemble prediction was equivalent to or greater than any individual model within it.

## 4. Discussion

Pathway-extended GE signatures generally improved accuracy of predicted patient responses to specific TKIs. Compared to signatures comprised solely of literature curated genes, PE signatures revealed previously unknown gene loci that contributed to drug response and, on average, had consistently better predictive performance. Aside from higher OA, the prediction accuracy for both sensitive and resistant patient groups (measured by MCC) was consistently more balanced. For example, **Cur-Lap** was the sole curated model with higher OA than its PE counterpart; however, its predictions were more skewed resulting in lower MCC. Furthermore, both MCC and overall accuracy were increased by AUC-weighted ensemble averaging of multiple PE models for sorafenib, erlotinib and imatinib. Except for lapatinib, the highest OAs were evident in patients receiving a ‘consensus’ prediction (where ≥75% of predictions made by the models in the ensemble predicted the same outcome for a patient). The improved predictive performance of PE SVMs, both individual and as ensembles of models, suggests that the genes within these signatures may refine the predominant mechanisms of both sensitivity and resistance to TKI therapy. PE gene models may be more useful in selecting chemosensitivity regimens for patients compared to models solely consisting of previously implicated genes known to respond to a specific chemotherapy.

Pathway-extension and the inclusion of pathway-related genes allowed for a larger pool of genes involved in ML. We avoided overfitting^78^ by pre-filtering these genes based on correlation with GI_50_. Furthermore, independent validation was determined by the identity and expression level of these features in patients treated with these drugs. Signatures containing pathway-related genes produced higher performing SVM signatures, consistent with the possibility that optimal molecular indicators of chemo-response may identify genes upstream or downstream of, or are interactors with, previously known cancer biomarkers. Generating SVMs from curated genes assures that features selected do not arise from statistical association alone. Generating PE SVMs required systematic selection of genes with established relationships to curated genes. For the best-performing PE gene signatures, most signature genes validated in the present study had been independently associated with abnormalities of expression, copy number or mutation in these tumour types (Additional References). Expanded signatures could potentially assist in the identification of novel biomarkers of chemo-response in these tissues.

Primary and secondary genes in PE gene signatures can offer context for drug responses without predicate literature support. The relationships between curated genes and genes selected through pathway-extension for sorafenib are illustrated in Figure 5. The vitamin D transporter encoded by *GC* is a major determinant of the response to this drug, as overall prediction accuracy is decreased by 52% upon its removal from **PE-Sor** (Figure 4A). In fact, *GC* is two nodes distant from multiple curated genes (*ABCB1*, *ABCC2* and *HNF1A*, among others [Figure 3A]). The *ABCB1* transporter has been implicated in sorafenib-related toxicities based on efflux efficiency.^79,80^ *ABCB1* also carries out efflux of Vitamin D_3_,^81^ and the 1,25-dihydroxy-vitamin D_3_ isoform (or 1.25D) activates *ABCB1* expression.^82^ Vitamin D is converted to this 1.25D isoform by *CYP27B1,* which is one-node distant from *ABCB1*. Similarly, *GC* binds specifically to 1.25D, which puts *GC* one node distant from *ABCB1*. The growth inhibitory effect of sorafenib has been shown to be amplified by 1.25D.^56^ Together, these network connections provide context that integrates functions and roles of individual genes of the tumour response to sorafenib. The PE signatures will be useful for understanding drug toxicity, although it was not explicitly a goal of this study. The importance of *GC* in **PE-Sor** may explain why a lower sorafenib dose is effective for treatment. Supplemental vitamin D_3_ reduces toxicity to sorafenib at this lower dose in differentiated thyroid carcinoma that is non-responsive to iodine therapy.^56^

The best performing SVMs for TKIs shared several common genetic pathways. Multiple PE models contained genes related to NOD-like receptor signaling (erlotinib: *NEK7*, *RELB*), PI3K-AKT signaling pathway (erlotinib: *CDK6*, *GNG3*; lapatinib: *ITGA11*; sunitinib: *EPHA2*, *NR4A1*) and Ras-Raf-MEK-ERK pathway (erlotinib: *CDK6*, *RELB*; sorafenib: *TGFB1*; sunitinib: *EPHA2*, *NR4A1*). Aberrant NOD-like receptor signaling drives carcinogenesis,^83^ while numerous cancer therapies target either or both of *PI3K* and *AKT*.^84–86^ The Ras-Raf-MEK-ERK pathway involves several protein kinases activated by tyrosine kinase receptors, with oncogenic mutations most prominently affecting Ras and B-Raf within the pathway.^87^ These pathways, which are disrupted broadly among different cancers, are implicated across numerous high performing ML models predicting TKI response.

Several pathway-extended (EPHA2, PRKD2, and PDGFRB) and curated (CDK6 and ABL2) gene products extrapolated from the highest performing signatures were bound to kinases based on a proteomic analysis of target selectivity for 243 kinase-inhibitors on 259 distinct tyrosine kinases.^88^ Few tyrosine kinase target genes from this proteomic analysis for TKIs in the current study exhibited correlations between GI_50_ and either gene expression or copy number (< 20° threshold; Table S7). *RET* was the only SVM gene implicated in the response to a TKI for both gene expression and protein (sorafenib; Concentration- and Target-Dependent Selectivity of 0.515; Klaeger et al. [2017]^88^). Therefore, expression of genes that are either positively or inversely correlated with drug response is generally unrelated to quantification of proteins that directly interact with the kinases themselves. If absence of signature genes from those corresponding to proteomic analysis is not attributable to either experimental or specific cell lines used, then signature gene expression is more likely indirectly regulated by gene products that are selective for most TKIs. Many genes in the PE SVMs were two nodes distant from curated biomarkers, which is consistent with the possibility that these represent common control points in the regulation of drug responses. In this regard, such control points exhibit behavior similar to state-cycle attractors of self-organizing systems.^89^ From a ML perspective, the dimensionality of the SVM model is reduced, avoiding overfitting, by substituting these control point genes for curated genes. Improvement in the prediction accuracy for both the sensitive and resistant patient categories might also be a consequence of these biomarkers being control points for *multiple* curated genes. Consider two curated genes that are “controlled” or regulated by the same two node biomarker, where inclusion of one of these improves accuracy for detecting drug sensitivity, and the other improves detection of resistance. Substituting the controlling gene for both curated genes in the PE-signature might improve accuracy of detection of both outcomes.

Transferability of these cell line-based models to other independent cell line datasets was also evaluated.^90^ PE TKI models were analyzed using data from the Sanger Genomics of Drug Sensitivity in Cancer Project (GDSC), including RNA-seq derived cancer cell line-derived gene expression data (E-MTAB-3983; ArrayExpress) based on IC_50_ values of cell lines in CancerRxGene.^91^ Using median IC_50_ to distinguish sensitivity from resistance, the top SVM that we derived for each TKI could not significantly separate cell lines sensitive and resistant to the same drug in GDSC (MCC from 0 to 0.19; OA ranging from 50-58%). Altering the IC_50_ thresholds did not significantly change these results. Applying this analysis to cell lines from specific tissues used in the derivation of the specific TKI signatures, **PE-Ima** was more accurate for seven imatinib-treated cell lines derived from intestinal tumours (OA of 69%; MCC – 0.41). The disparity in performance between the training and testing data sets may be related to differences in the expression patterns in different tissue types, or batch effects. IC_50_ measurements for the same cell line and drug are known to vary significantly between studies, especially when the cell line is drug insensitive,^92^ which may contribute to the poor correlation between results of both datasets.

Transferability of SVMs to different patient datasets may also be confounded by several other limitations of applying ML models derived from cell line expression to predict responses to the same drugs using patient GE data. By contrast with tumours, cancer cell lines tend to have a stable genetic profile when grown under controlled culturing conditions. Consequently, they tend to lack the genetic heterogeneity present in many tumour types,^93^ particularly during progression, which often occurs concomitant with evolution of acquired chemotherapy resistance.^94^ Cancer cell lines also lack extracellular matrix, which contributes to tumour growth, migration and invasion *in vivo*. These differences may challenge prediction accuracy of cell line-based SVMs using patient GE and/or CN. Clinical outcome measures within patient data sets were not consistent between different studies of the same tumour type. Finally, the cell line GE data used for training originated in this study solely from breast cancer, whereas patient tumour GE data were also derived from other cancer types.

## 5. Conclusions

The enhanced performance of chemotherapy response models developed using pathway-extension (over curated-only models) suggests that an interaction between a drug and its target may not directly relate with drug response; sensitivity could also be caused by a cellular event downstream of the drug-target interaction. PE models derived in this study demonstrated strong efficacy in selecting relevant genes, identifying novel molecular biomarker candidates, and predicting patient responses to TKIs. Strong-performing PE models appear to predict chemotherapy response in a cancer-type specific fashion, as many pathway-related genes selected by SVM software as novel candidate biomarkers of TKI efficacy were already prognostic biomarkers for the cancer type patients within the testing set were afflicted with. Ensemble averaging of multiple PE SVMs improved predictive accuracy in most cases and were found to be most commonly correct when predictions were highly consistent across each model constituting the ensemble. **PE-Erl** was also shown to have greater accuracy when considering solely female NSCLC patients. Interestingly, *RELB*, a feature in this signature, had previously demonstrated sexually dimorphic expression upon cancer treatment. The process of including pathway-related genes in biochemically inspired gene signatures can produce highly specific and accurate SVMs. PE models may have practical value, both in identifying novel biomarkers of chemosensitivity and in selecting effective chemotherapeutic agents.

## Supporting information

Additional References

## Acknowledgments

Compute Canada and Shared Hierarchical Academic Research Computing Network (SHARCNET) for a high-performance programming grant and computing facilities.

