## Supplementary figures and images for "Pathway-extended gene expression signatures integrate novel biomarkers that improve predictions of patient responses to kinase inhibitors"

### Figure S1

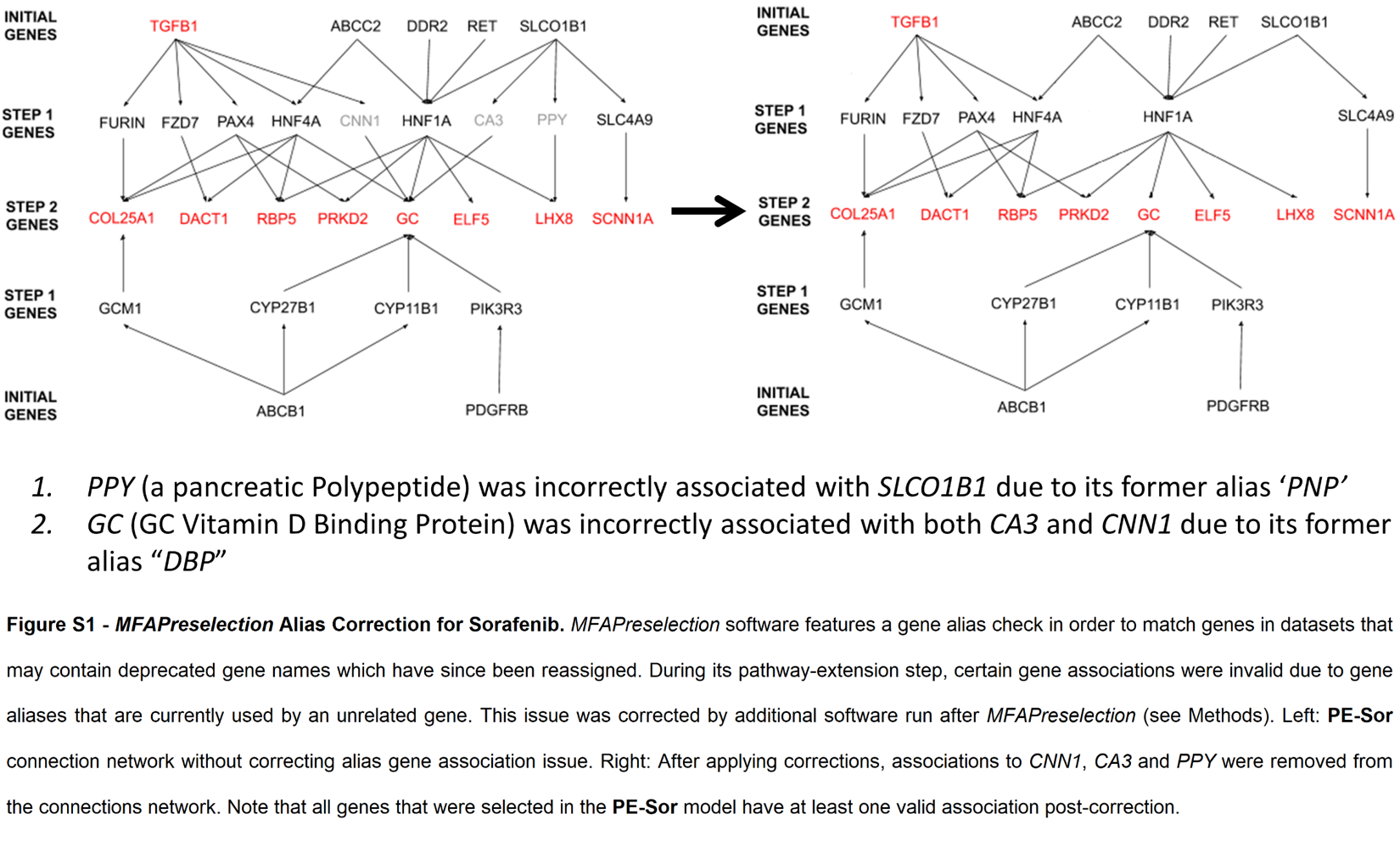

### Figure S2

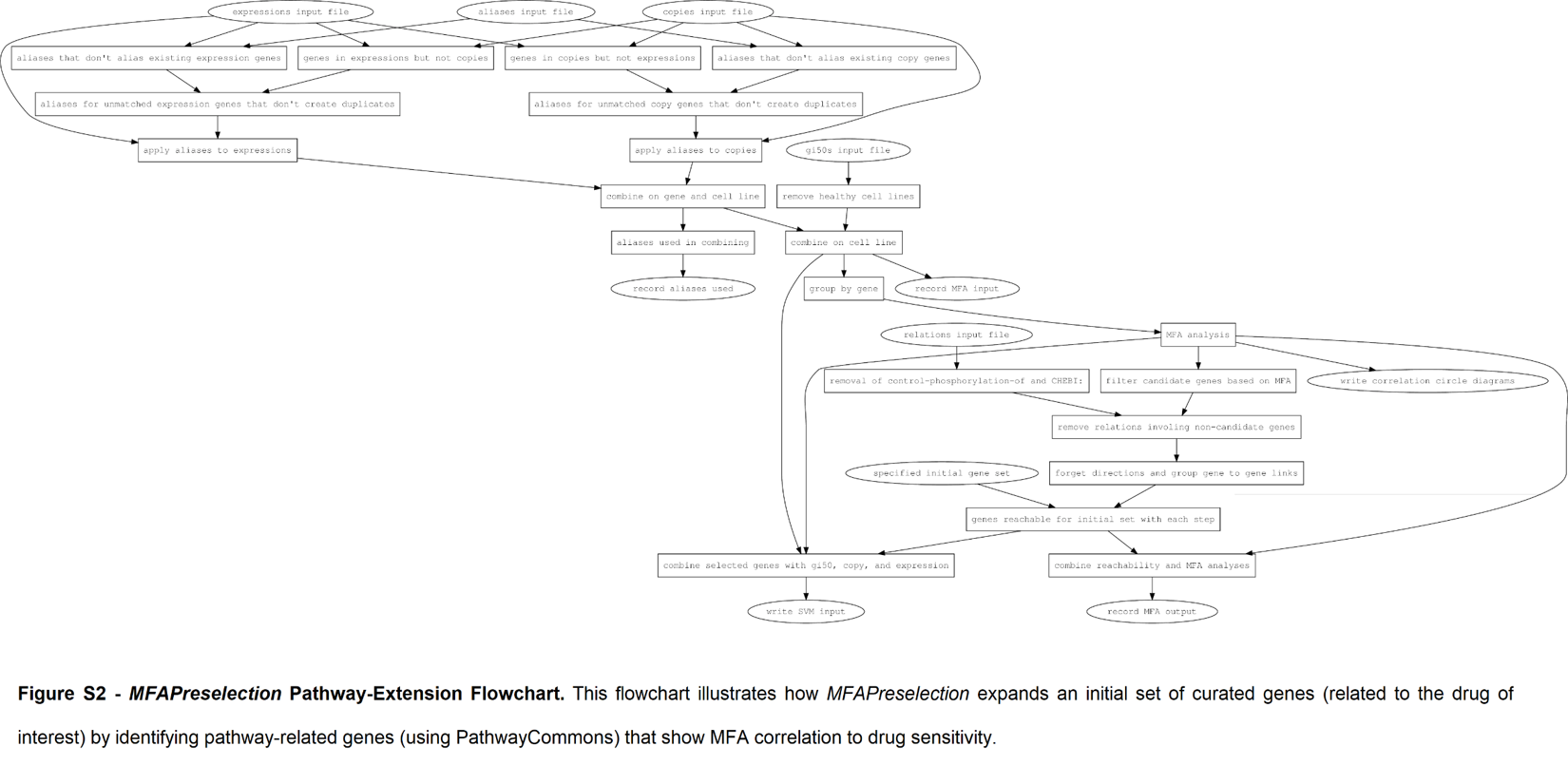
