## Additional References for "Pathway-extended gene expression signatures integrate novel biomarkers that improve predictions of patient responses to kinase inhibitors"

**Supplementary References**

**Pathway-extended multigene signatures improve predicted responses to kinase inhibitors by integrating novel biomarkers**

Ashis J. Bagchee-Clark^1^, Eliseos J. Mucaki^1^, Tyson Whitehead^2^, Peter K. Rogan^1,3*^

^1^ Department of Biochemistry, Schulich School of Medicine & Dentistry, University of

Western Ontario, Canada

^2^SHARCNET, London, Ontario, Canada

^3^ CytoGnomix Inc, London, Ontario, Canada

**Supplementary References.**

The following list indicates the studies that describe experimental evidence which support the inclusion of genes within the final signature and/or were involved in the generation of the final signature. We prioritized studies which showed evidence linking the gene to the pharmacokinetics, pharmacodynamics or efficacy of a TKI:

*CYP3A4 (Lapatinib)*

1. Towles JK, Clark RN, Wahlin MD, Uttamsingh V, Rettie AE, Jackson KD. Cytochrome P450 3A4 and CYP3A5-catalyzed bioactivation of lapatinib. *Drug Metabolism and Disposition.* 2016; 44(10), 1584-1597.

- In this study, *CYP3A4* was found to be an important contributor of lapatinib bioactivation.

*SNAI1 (Lapatinib)*

2. Desai K, Aiyappa R, Prabhu JS, et al. HR+ HER2− breast cancers with growth factor receptor–mediated EMT have a poor prognosis and lapatinib downregulates EMT in MCF-7 cells. *Tumor Biology.* 2017; 39(3), 1010428317695028.

- In this report, MCF7-cells treatment with lapatinib resulted in downregulation of epithelial-to-mesenchymal transition as indicated by lower levels of *SNAI1* transcripts (marker of epithelial-to-mesenchymal transition).

*INSR (Lapatinib)*

3. Zhang Z, Wang J, Ji D, et al. Functional genetic approach identifies MET, HER3, IGF1R, INSR pathways as determinants of lapatinib unresponsiveness in HER2-positive gastric cancer. *Clinical cancer research*. 2014; 20(17), 4559-4573.

- Here, the INSR pathway was identified as a lapatinib-resistant determinant in HER2-positive gastric cancer.

*ERBB2 (Lapatinib)*

4. Chu I, Blackwell K, Chen S, Slingerland J. The dual ErbB1/ErbB2 inhibitor, lapatinib (GW572016), cooperates with tamoxifen to inhibit both cell proliferation-and estrogen-dependent gene expression in antiestrogen-resistant breast cancer. *Cancer research*. 2005. 65(1), 18-25.

- In this study, lapatinib in combination with tamoxifen demonstrated efficacy treating breast cancer.

5. Iwata H, Fujii H, Masuda N, et al.. Efficacy, safety, pharmacokinetics and biomarker findings in patients with HER2-positive advanced or metastatic breast cancer treated with lapatinib in combination with capecitabine: results from 51 Japanese patients treated in a clinical study. *Breast Cancer.* 2015; 22(2), 192-200.

- Iwata et al reports significant improvement in treating ERBB2+ breast cancer patients upon lapatinib treatment in combination with capecitabine.

*BCL2L11 (Lapatinib)*

6. Park SH, Ito K, Olcott W, Katsyv I, Halstead-Nussloch G, Irie HY. PTK6 inhibition promotes apoptosis of Lapatinib-resistant Her2+ breast cancer cells by inducing Bim. *Breast Cancer Research*. 2015; 17(1), 86.

- In Park et al, induction of *BCL2L11* promoted apoptosis of lapatinib-resistant HER2+ breast cancer cells.

*NFKB1 (Lapatinib)*

7. Bailey ST, Miron PL, Choi YJ, et al. NF-κB activation-induced anti-apoptosis renders HER2-positive cells drug resistant and accelerates tumor growth. *Molecular Cancer Research*. 2014; 12(3), 408-420.

- This report details HER2 positive breast cancer cells rendered resistant to lapatinib via NFKB activation.

*NRG1 (Lapatinib)*

8. Leung WY, Roxanis I, Sheldon H, et al. Combining lapatinib and pertuzumab to overcome lapatinib resistance due to NRG1-mediated signalling in HER2-amplified breast cancer. *Oncotarget*. 2015; 6(8), 5678.

- Breast cancer cells were partially rescued via exogenous *NRG1* from lapatinib-induced growth inhibition.

*GRB7 (Lapatinib)*

9. Jernström S, Hongisto V, Leivonen SK, et al. Drug-screening and genomic analyses of HER2-positive breast cancer cell lines reveal predictors for treatment response. *Breast Cancer: Targets and Therapy*. 2017; 9, 185.

- Ten genes within this study, including *GRB7*, were found significantly associated with lapatinib response.

10. Nencioni A, Cea M, Garuti A, et al. Grb7 upregulation is a molecular adaptation to HER2 signaling inhibition due to removal of Akt-mediated gene repression. *PLoS One.* 2010; 5(2).

- Within this report, RNA-interference removal of *GRB7* was found to increase lapatinib activity.

*AXL (Lapatinib)*

11. Bouchalova K, Cizkova M, Cwiertka K, Trojanec R, Friedecky D, Hajduch M. Lapatinib in breast cancer-the predictive significance of HER1 (EGFR), HER2, PTEN and PIK3CA genes and lapatinib plasma level assessment. *Biomed Pap Med Fac Univ Palacky Olomouc Czech Repub.* 2010; 154(4), 281-288.

- In this manuscript, it is observed that *AXL*-overexpression is a characteristic of breast cancers that do not develop resistance to lapatinib.

*ABCB1 (Lapatinib)*

12. Dai CL, Tiwari AK, Wu CP, et al. Lapatinib (Tykerb, GW572016) reverses multidrug resistance in cancer cells by inhibiting the activity of ATP-binding cassette subfamily B member 1 and G member 2. *Cancer Research*. 2008; 68(19), 7905-7914.

- This paper details how *ABCB1* mediates drug resistance to lapatinib.

*TPD52 (Lapatinib)*

13. Parham LR, Briley LP, Li L, et al. Comprehensive genome-wide evaluation of lapatinib-induced liver injury yields a single genetic signal centered on known risk allele HLA-DRB1* 07: 01. *Pharmacogenomics J.* 2016; 16(2), 180-185.

- One intronic variant in *TPD52* (rs7828135) passed a pre-specified significance threshold in a genome-wide association study of lapatinib-induced liver injury, with respect to conditioning upon *HLA-DRB1*07:01* allele.

*MMP9 (Lapatinib)*

14. Umbreit C, Erben P, Faber A, et al. MMP9, Cyclin D1 and β-Catenin Are Useful Markers of p16-positive Squamous Cell Carcinoma in Therapeutic EGFR Inhibition In Vitro. *Anticancer Research*. 2015; 35(7), 3801-3810.

- Here, the *MMP9* expression profile is shown to be a potential early indicator of lapatinib sensitivity.

*TGFB1 (Sorafenib)*

15. Fernando J, Sancho P, Fernández‐Rodriguez CM, et al. Sorafenib sensitizes hepatocellular carcinoma cells to physiological apoptotic stimuli. *Journal of Cellular Physiology*. 2012; 227(4), 1319-1325.

- In this study, sorafenib sensitized hepatocellular carcinoma cells to the apoptotic activity of *TGFB1*.

*ABCC2 (Sorafenib)*

16. Wei D, Zhang H, Peng R, Huang C, Bai R. ABCC2 (1249G> A) polymorphism implicates altered transport activity for sorafenib. *Xenobiotica*. 2017; 47(11), 1008-1014.

- Wei et al demonstrates that *ABCC2* polymorphisms led to increases in MRP2 activity, resulting in increased sorafenib efflux.

*DDR2 (Sorafenib)*

17. Kumar R, Crouthamel MC, Rominger DH, Gontarek RR, Tummino PJ, Levin RA, King AG. Myelosuppression and kinase selectivity of multikinase angiogenesis inhibitors. *British Journal of Cancer*. 2009; 101(10), 1717-1723.

- *DDR2* was found to be inhibited by sorafenib.

*RET (Sorafenib)*

18. Plaza-Menacho I, Mologni L, Sala E, et al. Sorafenib functions to potently suppress RET tyrosine kinase activity by direct enzymatic inhibition and promoting RET lysosomal degradation independent of proteasomal targeting. *Journal of Biological Chemistry*. 2007; 282(40), 29230-29240.

- This paper reports that sorafenib directly inhibits RET activity.

*SLCO1B1 (Sorafenib)*

19. Zimmerman EI, Hu S, Roberts JL, et al. Contribution of OATP1B1 and OATP1B3 to the disposition of sorafenib and sorafenib-glucuronide. *Clinical Cancer Research*. 2013; 19(6), 1458-1466.

- This report outlines the involvement of *SLCO1B1* in sorafenib-glucuronide elimination in humanized transgenic mice.

*ABCB1 (Sorafenib)*

20. Qin C, Cao Q, Li P, et al. The influence of genetic variants of sorafenib on clinical outcomes and toxic effects in patients with advanced renal cell carcinoma. *Scientific Reports*. 2016; 6, 20089.

- Qin et al. finds that *ABCB1* polymorphisms lead to sorafenib-related toxicities in advanced renal cell carcinoma patients.

*PDGRFB (Sorafenib)*

21. Lierman E, Lahortiga I, Van Miegroet H, Mentens N, Marynen P, Cools J. The ability of sorafenib to inhibit oncogenic PDGFRβ and FLT3 mutants and overcome resistance to other small molecule inhibitors. *Haematologica*. 2007; 92(1), 27-34.

- Sorafenib inhibits ETV6-PDGFRB and FLT3 mutants.

*SHH (Gefitinib)*

22. Wang F, Wang W, Li J, Zhang J, Wang X, Wang M. Sulforaphane reverses gefitinib tolerance in human lung cancer cells via modulation of sonic hedgehog signaling. *Oncology Letters*. 2018; 15(1), 109-114.

- This paper details reversed gefitinib tolerance in human lung cancer cells through modulating *SHH* signaling via administration of sulforaphane.

*PLG (Gefitinib)*

23. Zhou J, Kwak KJ, Wu Z, et al. PLAUR confers resistance to Gefitinib through EGFR/P-AKT/Survivin signaling pathway. *Cellular Physiology and Biochemistry*. 2018; 47(5), 1909-1924.

- *PLG* encodes plasminogen, with knockdown of plasminogen activator reducing apoptosis of NSCLC cells.

*GCG (Gefitinib)*

24. Hare KJ, Hartmann B, Kissow H, Holst JJ, Poulsen SS. The intestinotrophic peptide, glp-2, counteracts intestinal atrophy in mice induced by the epidermal growth factor receptor inhibitor, gefitinib. *Clinical Cancer Research*. 2007; 13(17), 5170-5175.

- Simultaneous treatment with glucagon-like peptide-2 prevents small intestinal growth inhibition induced by gefitinib.

*CSF1 (Sunitinib)*

25. Chow LQ, Eckhardt SG. Sunitinib: from rational design to clinical efficacy. *Journal of Clinical Oncology*. 2007; 25(7), 884-896

- Sunitinib inhibits tyrosine kinases including colony-stimulating factor 1 (CSF-1).

*NFKB1 (Sunitinib)*

26. Zhu Y, Liu H, Xu L, et al. p21-activated kinase 1 determines stem-like phenotype and sunitinib resistance via NF-κ B/IL-6 activation in renal cell carcinoma. *Cell Death & Disease*. 2015; 6(2), e1637-e1637.

- Zhu et al. details *NFKB1* activation leading to sunitinib resistance in renal cell carcinoma cells.

*MET (Sunitinib)*

27. Zhou L, Liu XD, Sun M, et al. Targeting MET and AXL overcomes resistance to sunitinib therapy in renal cell carcinoma. *Oncogene*. 2016; 35(21), 2687-2697.

- This study reports that chronic sunitinib treatment induces activation of *MET* signaling.

*TSC1 (Sunitinib)*

28. Tran TA, Kinch L, Peña-Llopis S, Kockel L, Grishin N, Jiang H, Brugarolas J. Platelet-derived growth factor/vascular endothelial growth factor receptor inactivation by sunitinib results in Tsc1/Tsc2-dependent inhibition of TORC1. *Molecular and Cellular Biology*. 2013; 33(19), 3762-3779.

- Here, tumours with acquired *TSC1* mutations may be less responsive to sunitinib inhibition.

*VEGFC (Sunitinib)*

29. Dufies M, Giuliano S, Ambrosetti D, et al. Sunitinib stimulates expression of VEGFC by tumor cells and promotes lymphangiogenesis in clear cell renal cell carcinomas. *Cancer Research*. 2017; 77(5), 1212-1226.

- In this study, *VEGFC* in renal cell carcinoma cells is found to be stimulated by sunitinib.

*AXL (Sunitinib)*

30. Van Der Mijn JC, Broxterman HJ, Knol JC, et al. Sunitinib activates Axl signaling in renal cell cancer. *International Journal of Cancer*. 2016; 138(12), 3002-3010.

- Van Der Mijn et al finds that in renal cell carcinoma sunitinib activates *AXL*.

*ENPP2 (Sunitinib)*

31. Su SC, Hu X, Kenney PA, et al. Autotaxin–lysophosphatidic acid signaling axis mediates tumorigenesis and development of acquired resistance to sunitinib in renal cell carcinoma. *Clinical Cancer Research*. 2013; 19(23), 6461-6472.

- This paper finds endothelial *ENPP2* is involved in acquiring resistance to sunitinib.

*FGB (Sunitinib)*

32. Dzik C, Reis ST, Viana NI, et al. Gene expression profile of renal cell carcinomas after neoadjuvant treatment with sunitinib: New pathways revealed. *Int J Biol Markers*. 2017. 32(2), 210-217.

- Study measured lower expression levels of *FGB* after sunitinib treatment compared with controls.

*RUNX1 (Imatinib)*

33. Miething C, Grundler R, Mugler C, et al. Retroviral insertional mutagenesis identifies RUNX genes involved in chronic myeloid leukemia disease persistence under imatinib treatment. *Proceedings of the National Academy of Sciences*. 2007; 104(11), 4594-4599.

- Stable or inducible expression of *RUNX1* in Bcr-Abl-positive cell lines led to protection from imatinib-induced apoptosis.

*RET (Imatinib)*

34. de Groot JWB, Menacho IP, Schepers H, et al. Cellular effects of imatinib on medullary thyroid cancer cells harboring multiple endocrine neoplasia Type 2A and 2B associated RET mutations. *Surgery*. 2006; 139(6), 806-814.

- This manuscript finds imatinib inhibits *RET*-mediated medullary thyroid carcinomas cells in a dose-dependent manner.

*IGF1R (Imatinib)*

35. Martins AS, Mackintosh C, Martín DH, Campos M, Hernández T, Ordóñez JL, de Alava E. Insulin-like growth factor I receptor pathway inhibition by ADW742, alone or in combination with imatinib, doxorubicin, or vincristine, is a novel therapeutic approach in Ewing tumor. *Clinical Cancer Research*. 2006; 12(11), 3532-3540.

- Martins et al. demonstrates that addition of imatinib to ADW742, an IGF-1R inhibitor, augmented apoptosis of Ewing tumor cell lines with high *IGF1R* activation levels.

*ABCC2 (Imatinib)*

36. He B, Bai Y, Kang W, Zhang X, Jiang X. LncRNA SNHG5 regulates imatinib resistance in chronic myeloid leukemia via acting as a CeRNA against MiR-205-5p. *American Journal of Cancer Research*. 2017; 7(8), 1704.

- Here, imatinib resistance in chronic myeloid leukemia cells is promoted via regulation of *ABCC2* by lncRNA SNHG5.

37. Au A, Baba AA, Azlan H, Norsa'adah B, Ankathil, R. Clinical impact of ABCC 1 and ABCC 2 genotypes and haplotypes in mediating imatinib resistance among chronic myeloid leukaemia patients. *Journal of Clinical Pharmacy and Therapeutics*. 2014; 39(6), 685-690.

- This paper details three polymorphisms in *ABCC2* (‐24C>T, 1249G>A and 3972C>T) creating a haplotype associated with imatinib response.

*CDKN1A (Imatinib)*

38. Ferrandiz N, Caraballo JM, Albajar M, et al. p21Cip1 Confers resistance to imatinib in human chronic myeloid leukemia cells. *Cancer Letters*. 2010; 292(1), 133-139.

- In this study, p21 (*CDKN1A*) is found to confer resistance to imatinib in human chronic myeloid leukemia cells.

*MYB (Imatinib)*

39. Ohmine K, Nagai T, Tarumoto T, et al. Analysis of gene expression profiles in an imatinib‐resistant cell line, KCL22/SR. *Stem Cells*. 2003; 21(3), 315-321.

- Here, c-MYB was downregulated in KCL22/SR chronic myelogenous leukemia cells resistant to imatinib.

*BCL2 (Imatinib)*

40. Ko TK, Chuah CT, Huang JW, Ng KP, Ong ST. The BCL2 inhibitor ABT-199 significantly enhances imatinib-induced cell death in chronic myeloid leukemia progenitors. *Oncotarget*. 2014; 5(19), 9033.

- This study defines how, in chronic myeloid leukemia cells, administration of *BCL2*-inhibitor ABT-199 significantly enhances imatinib-induced cell death.

*PDGFRB (Imatinib)*

41. Cheah CY, Burbury K, Apperley JF, et al. Patients with myeloid malignancies bearing PDGFRB fusion genes achieve durable long-term remissions with imatinib. *Blood.* 2014; 123(23), 3574-3577.

- Here, imatinib was found to achieve excellent long-term responses in patients with myeloid malignancies bearing *PDGFRB* fusion genes.

*GZMB (Imatinib)*

42. Stacchiotti S, Pantaleo MA, Negri T, et al. Efficacy and biological activity of imatinib in metastatic dermatofibrosarcoma protuberans (DFSP). *Clinical Cancer Research*. 2016; 22(4), 837-846.

- *GZMB* was upregulated in imatinib-treated dermatofibrosarcoma protuberans/fibrosarcomatous samples.

*PTGS2 (Imatinib)*

43. Johnson FM, Yang P, Newman RA, Donato NJ. Cyclooxygenase-2 induction and prostaglandin E 2 accumulation in squamous cell carcinoma as a consequence of epidermal growth factor receptor activation by imatinib mesylate. *J Exp Ther Oncol*. 2004;4(4):317-25.

- In this study, COX-2 (*PTGS2*) expression was induced in an imatinib dose-dependent manner.

*IL3 (Imatinib)*

44. Abrishami M, Pastushok L, Mochoruk K, DeCoteau JF, Geyer CR. Synthetic Anti-IL3 Receptor Antibodies As Therapeutics to Block Innate Imatinib Resistance in Chronic Myelogenous Leukemia. *Blood*. 2014; 124(21):4519.

- IL-3 presence reduced apoptosis induction and cytotoxicity by imatinib.

*PTK2B (Imatinib)*

45. Ovcharenko A, Granot G, Rokah OH, Park J, Shpilberg O, Raanani P. Enhanced adhesion/migration and induction of Pyk2 expression in K562 cells following imatinib exposure. *Leukemia Research*. 2013; 37(12), 1729-1736.

- In this paper, Pyk2 mRNA and protein levels were induced by imatinib administration.

*CYP2C19 (Imatinib)*

46. Sacha T. (2014). Imatinib in chronic myeloid leukemia: an overview. *Mediterr J Hematol Infect Dis*. 2014;6(1):e2014007.

- *CYP2C19* contributes to imatinib metabolism.

*CYP2C9 (Imatinib)*

46. Sacha T. (2014). Imatinib in chronic myeloid leukemia: an overview. *Mediterr J Hematol Infect Dis*. 2014;6(1):e2014007.

- *CYP2C9* contributes to imatinib metabolism.

*MET (Erlotinib)*

47. Costa DB, Nguyen KSH, Cho BC, et al. Effects of erlotinib in EGFR mutated non-small cell lung cancers with resistance to gefitinib. *Clinical Cancer Research*. 2008; 14(21), 7060-7067.

- *MET* amplification is a mechanism of gefitinib resistance *in vitro.*

*IGF1R (Erlotinib)*

48. Hussmann D, Madsen AT, Jakobsen KR, Luo Y, Sorensen BS, Nielsen AL. IGF1R depletion facilitates MET-amplification as mechanism of acquired resistance to erlotinib in HCC827 NSCLC cells. *Oncotarget*. 2017; 8(20), 33300.

- Here, acquired resistance of erlotinib in NSCLC cells via MET-amplification is facilitated by *IGF1R* depletion.

*BAX (Erlotinib)*

49. Ling YH, Lin R, Perez-Soler R. RETRACTION: Erlotinib Induces Mitochondrial-Mediated Apoptosis in Human H3255 Non-Small-Cell Lung Cancer Cells with Epidermal Growth Factor ReceptorL858R Mutation through Mitochondrial Oxidative Phosphorylation-Dependent Activation of BAX and BAK. *Molecular Pharmacology*. 2008; 74(3), 793-806.

- Erlotinib-induced apoptosis was reduced after down-regulation of *BAX* gene expression.

*CDK6 (Erlotinib)*

50. Zhou J, Wu Z, Wong G, et al. CDK4/6 or MAPK blockade enhances efficacy of EGFR inhibition in oesophageal squamous cell carcinoma. *Nature Communications*. 2017; 8(1), 1-12.

- In this study, erlotinib response in esophageal squamous cell carcinoma xenografts was improved via inhibition of *CDK4/6*.

*TGFB3 (Erlotinib)*

51. Halatsch ME, Löw S, Mursch K, et al. Candidate genes for sensitivity and resistance of human glioblastoma multiforme cell lines to erlotinib. *Journal of Neurosurgery*. 2009; 111(2), 211-218.

- This study finds, in glioblastoma multiforme cell lines, *TGFB3* was associated with resistance to erlotinib.

*FOXO1 (Erlotinib)*

52. Chen X, Zhu L, Ma Z, et al. Oncogenic miR-9 is a target of erlotinib in NSCLCs. *Scientific Reports*. 2015; 5, 17031.

- FOXO1 protein expression was upregulated by erlotinib.
